# Subunit-dependent and independent rules of AMPA receptor trafficking during long-term depression in hippocampal neurons

**DOI:** 10.1101/2020.11.12.379867

**Authors:** Shinji Matsuda, Michisuke Yuzaki

## Abstract

Long-term potentiation (LTP) and depression (LTD) of excitatory neurotransmission are believed to be the neuronal basis of learning and memory. Both processes are primarily mediated by neuronal activity-induced transport of postsynaptic AMPA-type glutamate receptors (AMPARs). While AMPAR subunits and their specific phosphorylation sites mediate differential AMPAR trafficking, LTP and LTD could also occur in a subunit-independent manner. Thus, it remains unclear whether and how, certain AMPAR subunits with phosphorylation sites are preferentially recruited to or removed from synapses during LTP and LTD. Here, we show that phosphorylation of the membrane-proximal region (MPR), which only occurs in GluA1 AMPAR subunits, mediates the subunit-dependent endosomal transport of AMPARs during LTD. AP-2 and AP-3, adaptor protein complexes necessary for clathrin-mediated endocytosis and late endosomal/lysosomal trafficking, respectively, are reported to be recruited to AMPARs by binding to the AMPAR auxiliary subunit, stargazin (STG), in an AMPAR subunit-independent manner. However, the association of AP-3, but not AP-2, with STG was indirectly inhibited by the phosphomimetic mutation in the MPR of GluA1. Thus, although AMPARs containing the phosphomimetic mutation at the MPR of GluA1 were endocytosed by a LTD-inducing stimulus, they were quickly recycled back to the cell surface in hippocampal neurons. These results could explain how the phosphorylation status of GluA1-MPR plays a dominant role in subunit-independent STG-mediated AMPAR trafficking during LTD.

## Introduction

Long-term potentiation (LTP) and long-term depression (LTD) of excitatory neurotransmission at glutamatergic synapses have been intensively studied as the neural basis of learning and memory (1,2). LTP and LTD are mainly caused by changes in the number of postsynaptic AMPA-type glutamate receptors (AMPARs) through activity-dependent lateral diffusion of AMPARs from or to postsynaptic sites, coupled with endosomal transport of AMPARs by exocytosis or endocytosis (3,4). GluA1 and GluA4 AMPAR subunits are primarily recruited to synapses in an activity-dependent manner (5) during LTP. In contrast, N-methyl-D-aspartate receptor (NMDAR) activation was shown to preferentially induce endocytosis of GluA2-containing AMPARs, followed by subsequent transport to the late endosome/lysosome pathway during LTD (6). In contrast, GluA2-lacking AMPARs are recycled back to the cell surface (6). Indeed, LTD is impaired in the cerebellum lacking GluA2 expression (7). Furthermore, phosphorylation of the GluA1 C-terminus by calcium/calmodulin-dependent protein kinase II (CaMKII; Ser831) and protein kinase A (PKA; Ser 845) have been shown to regulate LTP and LTD (8,9). Phosphorylation at Ser818 by protein kinase C (PKC) and phosphomimetic mutation at Ser 816 were shown to promote synaptic incorporation of GluA1 (10,11) (Figure 1A). These findings indicate that activity-dependent AMPAR trafficking is determined by the C-terminus of GluA subunits. However, such subunit-specific “rules” have been challenged by recent findings that LTP (12) and LTD (13) do not require the C-termini of GluA subunits.

**Figure 1.**
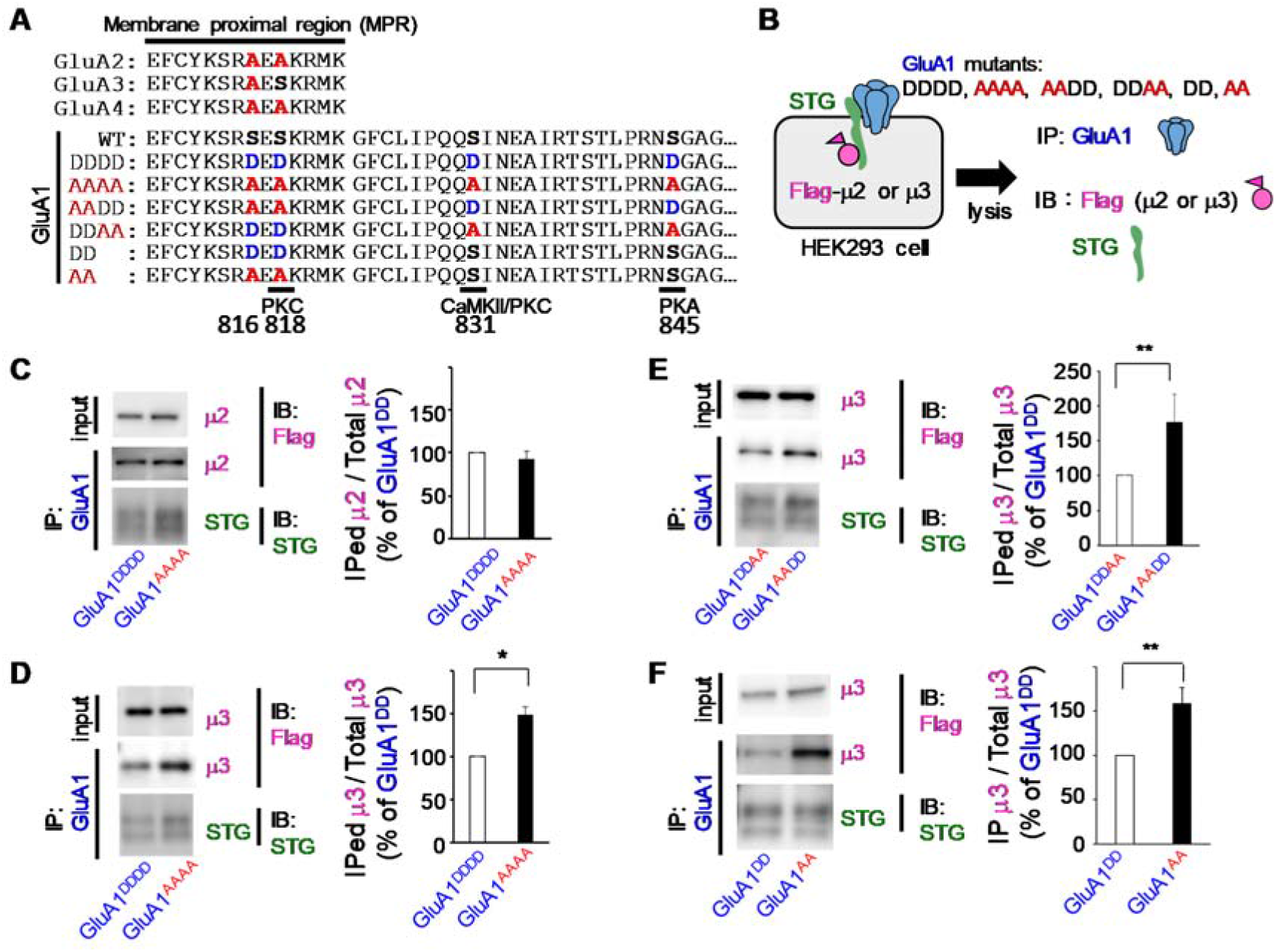
Phosphorylation of MPR regulates the affinity of the AMPA receptor-TARP complex to AP-3. **A.** Amino acid sequences of the C-terminus of AMPAR subunits and GluA1 mutants. Serine residues that can be phosphorylated by PKC, CaMKII, and PKA are indicated. These residues were replaced with aspartate and alanine to mimic phosphorylation (blue) and dephosphorylation (red). Although Ser816 is not directly phosphorylated, it enhances the effect of the Ser818 mutation. **B**. Schematic drawing of the co-immunoprecipitation assay. Lysates of HEK293 cells expressing STG, GluA1 mutants, and flag-tagged μ2 or μ3 were immunoprecipitated using the anti-GluA1 antibody. **C, D.** The effect of mutation of all serine residues of GluA1 on the interaction with μ2 or μ3. While μ2 was similarly co-immunoprecipitated with GluA1^AAAA^ and GluA1^DDDD^ (**C**), μ3 was preferentially co-immunoprecipitated with GluA1^AAAA^ than GluA1^DDDD^ (**D**). The intensity of the band corresponding to μ2 or μ3 that was co-immunoprecipitated was normalized to the intensity ofthe respective molecule in the input lysate. Data are presented as mean + SEM (Mann-Whitney U-test, *p < 0.05; *n* = 4). **E.** The effect of the position of the mutations on the interaction with μ3. μ3 was preferentially co-immunoprecipitated with GluA1^AADD^ than with GluA1^DDAA^. The intensity of μ3 in the immunoprecipitated fraction was normalized to that of the input lysate. Data are presented as mean + SEM (Mann-Whitney U-test, **p < 0.01; *n* = 6). **F.** The effect of mutations in the MPR on the interaction with μ3. μ3 was preferentially co-immunoprecipitated with GluA1^AA^ than with GluA1^DD^. The intensity of μ3 in the immunoprecipitated fraction was normalized to that of the input lysate. Data are presented as mean + SEM (Mann-Whitney U-test, **p < 0.01; *n* = 5).

An alternative hypothesis is that AMPAR trafficking is regulated by its auxiliary subunits, such as transmembrane AMPAR regulatory proteins (TARPs), which bind to all AMPAR subunits indiscriminately. The C termini of TARPs stabilize postsynaptic AMPARs by binding to anchoring proteins, such as postsynaptic density 95 (PSD95) (14). The C-terminus of TARPs contains multiple phosphorylation sites for CaMKII, PKC, and PKA, and positively charged residues (15). Phosphorylation of the C-termini of TARPs enhances its binding to PSD95 by neutralizing positive charges (16). Furthermore, we previously showed that dephosphorylated TARPs could specifically interact with the μ subunit of the adaptor proteins AP-2 (μ2) and AP-3 (μ3), which are essential for clathrin-dependent endocytosis and late endosomal/lysosomal trafficking, respectively (17). Thus, activity-dependent phosphorylation status of TARPs during LTP/LTD could affect lateral diffusion of postsynaptic AMPARs, followed by their endocytosis, in a manner independent of AMPAR subunits.

Recently, using mouse lines in which the endogenous C-termini of GluA1 and GluA2 were replaced with each other, the C-termini of GluA1 and GluA2 were shown to be necessary and sufficient for hippocampal LTP and LTD, respectively (18). Thus, we hypothesized that AMPAR subunits and their phosphorylation status were mechanistically linked with TARP-mediated trafficking. In the present study, we examined whether and how the phosphorylation of GluA1 C-terminus could affect its association with STG, a prototype of TARP, and adaptor proteins AP-2 and AP-3. We show that although the PKC phosphorylation sites of GluA1 do not affect its interaction with STG, phosphorylation of GluA1indirectly inhibits AP-3 binding to STG. Unless GluA1 was fully dephosphorylated, NMDA-induced LTD was impaired in hippocampal neurons, indicating that TARP-mediated AMPAR trafficking was affected by a subunit-specific rule.

## Results

### Phosphorylation of GluA1-MPR affects AP-3 binding to STG

The C-terminus of GluA1, but not GluA2, contains three serine residues that can undergo phosphorylation by PKC, CaMKII, and PKA (3,19) (Figure 1A). To test the hypothesis that the phosphorylation status of GluA1 may affect TARP-mediated AMPAR trafficking, we replaced all four serine residues with aspartate (GluA1^DDDD^) and alanine (GluA1^AAAA^), to mimic phosphorylated and dephosphorylated GluA1, respectively. We co-expressed GluA1 mutants, STG, and flag-tagged μ2 or μ3 subunits in human embryonic kidney 293 (HEK293) cells and performed co-immunoprecipitation assays (Figure 1B). Although GluA1^DDDD^ and GluA1^AAAA^ similarly co-immunoprecipitated STG, the amount of μ3, but not μ2, that co-immunoprecipitated with GluA1^DDDD^ was lower than that with GluA1^AAAA^ (Figure 1 C and D). To determine the serine residues responsible, we generated GluA1^AADD^ and GluA1^DDAA^, in which either S816/S818 or S831/S845 were replaced with alanine or aspartate without changing the total number of phosphomimetic sites (Figure 1A). Although GluA1^AADD^ and GluA1^DDAA^ similarly co-immunoprecipitated STG, the amount of μ3 that was co-immunoprecipitated by GluA1^DDAA^ was lower than that by GluA1^AADD^ (Figure 1 E), indicating that phosphorylation at Ser816/Ser818 likely reduced the interaction of STG with μ3. Indeed, GluA1^DD^, in which Ser816/Ser818 was replaced with aspartate, co-immunoprecipitated a significantly smaller amount of μ3 than GluA1^AA^, in which S816/S818 were replaced with alanine (Figure 1F, *n* = 5, p = 0.008 by Mann-Whitney *U*-test). These results indicate that phosphorylation at the membrane-proximal region (MPR) of GluA1 (Figure 1A) affects AP-3 binding to the AMPAR-STG complex.

### GluA1-MPR enhances the interaction between STG and AP-3

To assess how the MPR of GluA1 affects interaction of μ3 with STG, we prepared the C-terminus of STG as a glutathione S-transferase (GST) fusion protein and performed pull-down assays using cell lysates of HEK293 cells expressing flag-tagged μ2 or μ3. We synthesized the MPR peptide mimicking phosphorylated (MPR^DD^) or un-phosphorylated (MPR^AA^) GluA1 and added it to the lysate (Figure 2 A). The presence of MPR^AA^ or MPR^DD^ did not affect the amount of μ2 pulled-down by GST-STG (Figure 2B, p = 0.99 by Kruskal-Wallis test). Interestingly, the amount of μ3 pulled down by GST-STG was significantly increased by addition of the MPR^AA^ peptide (Figure 2 C, MPR^AA^, 126 ± 18%; MPR^DD^, 100 %; without MPR, 86 ± 13%; p = 0.006, MPR^AA^ vs. MPR^DD^; p = 0.043, MPR^AA^ vs. −MPR, *n* = 6 each by Kruskal-Wallis test and Steel-Dwass post hoc test). These results indicate that the presence of unphosphorylated MPR of GluA1 selectively enhanced the interaction between STG and AP-3.

**Figure 2.**
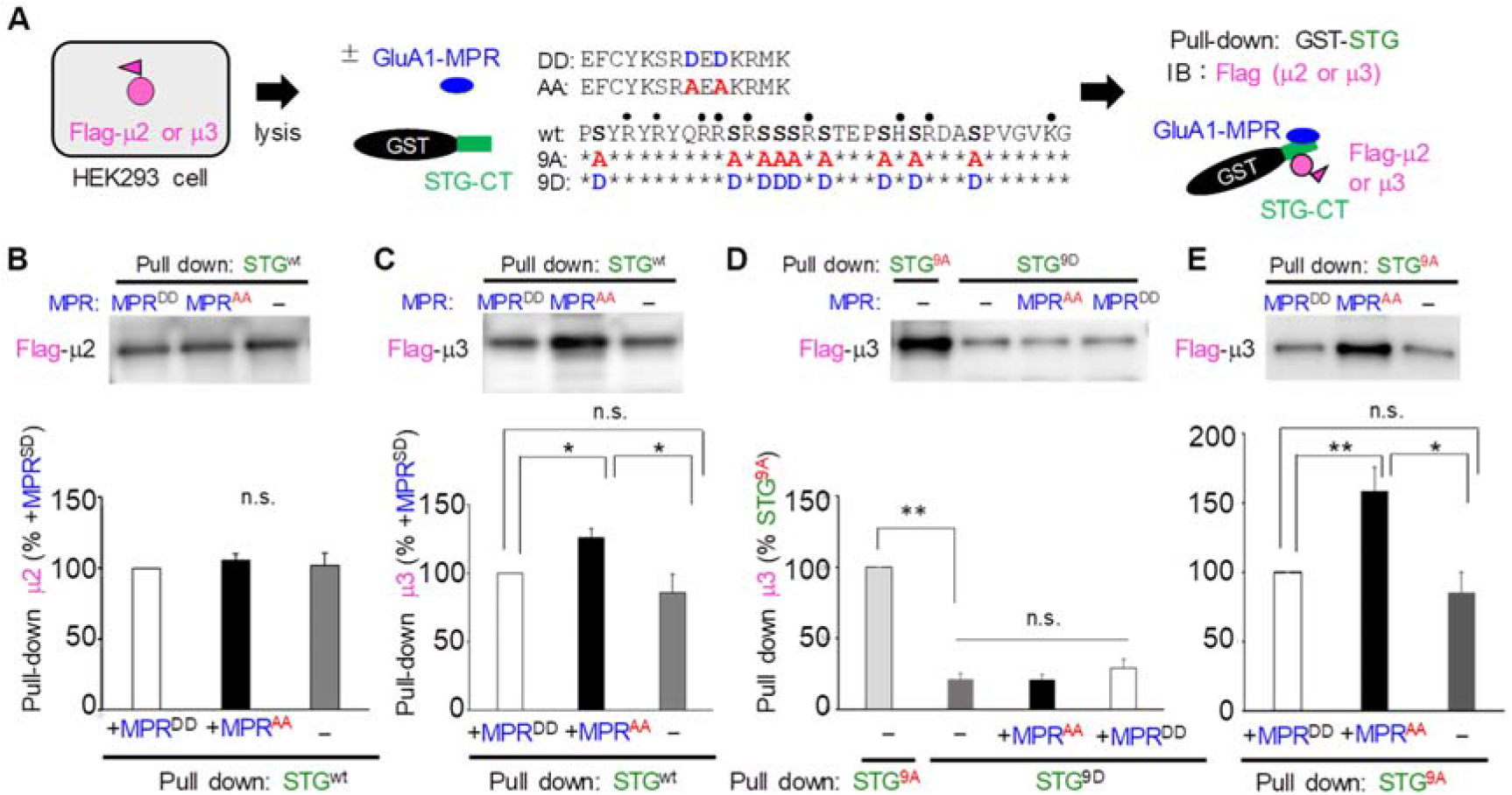
Dephosphorylated MPR enhances the interaction between STG and AP-3. **A.** Schematic drawing of the pull-down assay. Lysates of HEK293 cells expressing flag-tagged μ2 or μ3 were pulled down with the GST-fused C-terminus of STG (GST-CT) in the presence or absence of synthetic peptides corresponding to the MPR of GluA1. Amino acid sequences of the MPR and STG-CT, in which serine residues were replaced with alanine (red) or aspartate (blue) to mimic phosphorylated and dephosphorylated forms, are shown. **B, C.** Pull-down assays showing the effect of MPR on the interaction between wild-type STG and μ2 or μ3. The amount of μ2 or μ3 that was pulled down with GST-STG^wt^ in the presence of MPR^DD^ was arbitrarily established as 100%. The addition of MPR^DD^ or MPR^AA^ did not affect the interaction between STG^wt^ and μ2 (**B**), whereas MPR^AA^ enhanced the interaction between STG^wt^ and μ3 (**C**). Data are presented as mean + SEM. Kruskal-Wallis test and Steel-Dwass post hoc test, *p < 0.05; n = 6 each. **D.** Pull-down assays showing the effect of MPR on the interaction between μ3 and STG^9A^ or STG^9D^. The amount of μ3 that was pulled down with GST-STG^9A^ without the addition of MPR was arbitrarily established as 100%. Phosphomimetic mutation of STG (STG^9D^) significantly reduced the amount of pulled down μ3. Data are presented as mean + SEM. Mann-Whitney U-test, **p < 0.01; *n* = 6 each. The MPR peptides did not affect the interaction between μ3 and STG^9D^. Kruskal-Wallis test and Steel-Dwass post hoc test, *n* = 6 each. **E.** Pull-down assays showing the effect of MPR on the interaction between STG^9A^ and μ3. The amount of μ3 that pulled down with GST-STG^9A^ in the presence of MPR^DD^ was arbitrarily established as 100%. The addition of MPR^AA^ enhanced the interaction between STG^9A^ and μ3. Data are presented as mean + SEM. Kruskal-Wallis test and Steel-Dwass post hoc test, **p < 0.01, *p < 0.05; *n* = 6 each.

STG itself contains multiple positively charged residues and phosphorylation sites at the C-terminus (Figure 2 A). We next examined whether the facilitatory effect of MPR^AA^ on STG-AP-3 interaction was affected by the phosphorylation status of STG. As reported previously, the amount of μ3 pulled down by GST-STG^9D^, in which nine serine residues were replaced with aspartate to mimic phosphorylated STG, was significantly lower than that pulled down by GST-STG^9A^, mimicking the unphosphorylated form (Figure 2 D, STG^9A^, 100%; STG^9D^, 21 ± 3%; p = 0.002, *n* = 6 each, by Mann-Whitney *U*-test). The presence of MPR^AA^ or MPR^DD^ did not affect the amount of μ3 pulled down by STG^9D^ (Figure 2 D, p = 0.48 Kruskal-Wallis test). In contrast, the amount of μ3 pulled down by STG^9A^ was significantly increased by the addition of MPR^AA^ (Figure 2 E). These results indicate that dephosphorylation is required for STG to bind μ3 and that the presence of unphosphorylated GluA1-MPR further enhances STG-μ3 association.

### GluA1-MPR directly binds STG and indirectly enhances STG-AP3 interaction

To examine whether and how the MPR of GluA1 binds to the C-terminus of STG, we synthesized biotinylated MPR^DD^ and MPR^AA^ and performed a pull-down assay using streptavidin beads (Figure 3 A). GluA1-MPR^AA^ pulled down GST-STG much more than MPR^DD^ (Figure 3 B). To identify the region of STG necessary for MPR binding, we prepared GST-STG^CT1^ and GST-STG^CT12^, in which the C-terminus of STG was sequentially deleted (Figure 3C). Although STG^wt^ and STG^CT12^ were similarly pulled down by GluA1-MPR^AA^, STG^CT1^ was not (Figure 3D), indicating that the CT2 region (230–259) was mediating binding to the MPR of GluA1. When the cell lysates from HEK293 cells expressing flag-tagged μ3, were pulled down by biotinylated MPR^AA^ in the presence of GST or GST-STG^wt^ (Figure 3 A), a large amount of μ3 was pulled down by MPR^AA^ in the presence of GST-STG^wt^ compared with GST (Figure 3E, GST only, 100%; GST-STG^wt^, 180 ± 34%; p = 0.0003, by Mann-Whitney U test), indicating that μ3 indirectly associates with the STG-MPR complex. Together, we propose that dephosphorylated STG directly binds to AP-3, and that dephosphorylated GluA1-MPR could further bind to STG and indirectly enhance the GluA1-STG complex (Figure 3F).

**Figure 3.**
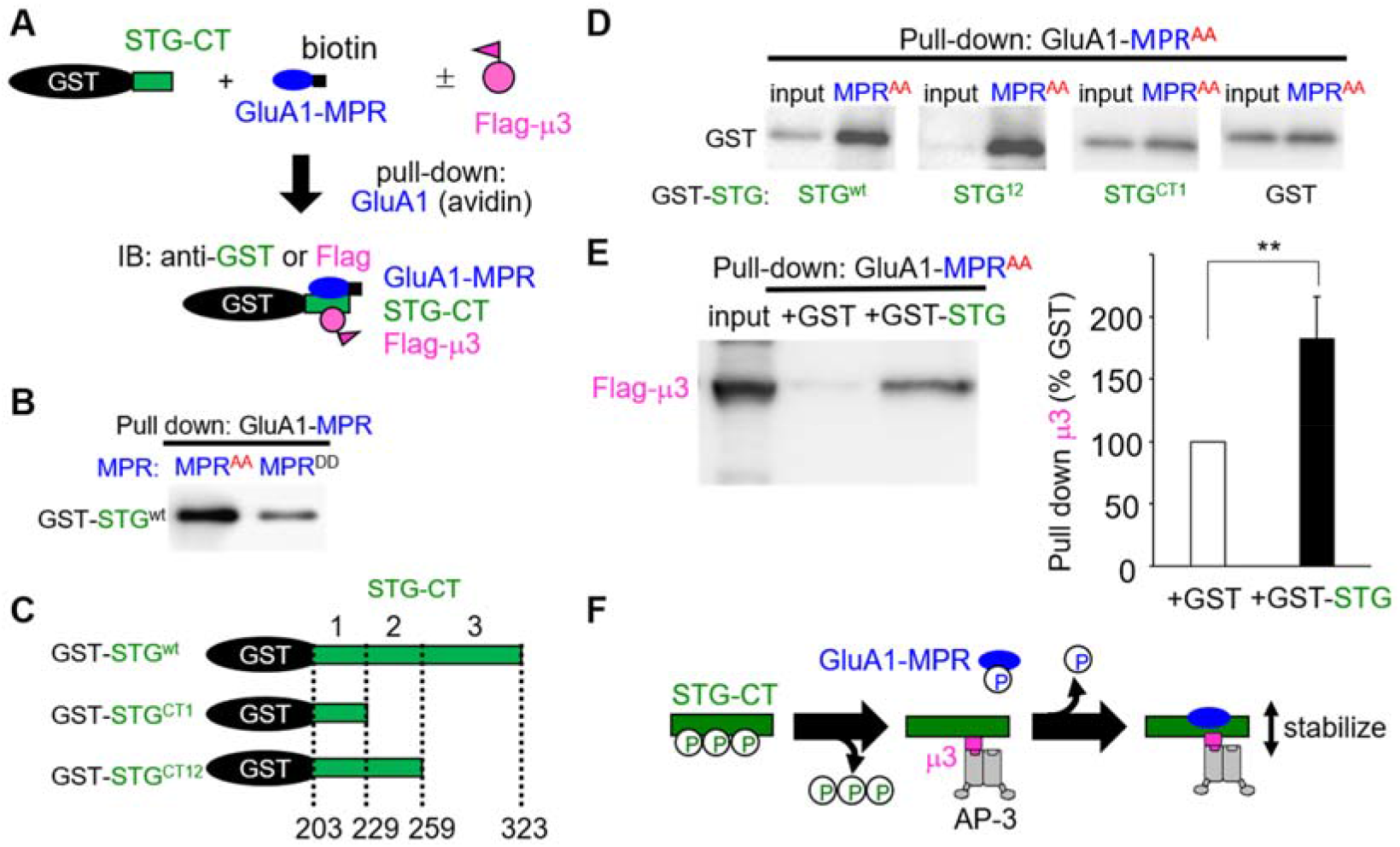
Dephosphorylated MPR directly binds to the C-terminus of STG. **A.** Schematic drawing of the pull-down assay. GST-fused C-terminus of STG (STG-CT) was pulled down using avidin that interacted with a synthetic biotinylated MPR peptide. In some experiments, lysates of HEK293 cells expressing flag-tagged μ3 were added. **B.** Pull-down assays showing a direct interaction between STG-CT and MPR. Dephosphomimetic MPR (MPR^AA^) showed a stronger interaction with STG-CT than phosphomimetic MPR (MPR^DD^). **C.** Schematic drawing of the deletion mutants of the GST-fused C-terminus of STG. Lower numbers indicate the amino acid position of full-length STG. **D.** Pull-down assays showing the interaction between STG deletion mutants and GluA1-MPR. The amount of STG pulled down with GluA1-MPR^AA^ was reduced by the deletion of amino acids 229–259 (STG^CT1^). **E**. Pull-down assays showing GluA1-MPR indirectly associates with μ3 via STG. A larger amount of μ3 was pulled down by GluA-MPR^AA^ when lysates of HEK293 cells expressing flag-tagged μ3 were added. The amount of μ3 pulled down with MPR^AA^ in the presence of GST was arbitrary established as 100%. Data are presented as mean + SEM. Mann-Whitney U-test, **p < 0.01; n = 8. **F.** Schematic drawing of the enhanced interaction between STG and AP-3 by addition of dephosphorylated MPR. Dephosphorylated STG can interact with AP-3, and this interaction is further enhanced by the binding of dephosphorylated GluA1-MPR to the CT2 region of STG.

### Phosphorylation of GluA1-MPR regulates NMDA-induced LTD

To clarify the role of phosphorylation of GluA1-MPR on AMPAR trafficking, we used a chemical LTD model, in which NMDA application induces AMPAR endocytosis (6,17). We expressed mutant GluA1, in which a hemagglutinin (HA) tag was added to the N-terminal extracellular domain, and Ser816/Ser818 were replaced with aspartate (GluA1^DD^) or alanine (GluA1^AA^), in cultured hippocampal neurons. After treatment with NMDA (50 μM) for 10 min, the cell surface and total GluA1 were sequentially detected by an anti-HA antibody before and after permeabilizing the plasma membrane (Figure 4 A). The intensity of cell surface HA-GluA1^AA^ was significantly reduced by NMDA treatment (control, 100 ± 10%, *n* = 13 cells; NMDA, 71 ± 8%; *n* = 14 cells; p = 0.033 by two-tailed Student’s *t*-test; Figure 4 B). In contrast, the intensity of cell surface HA-GluA1^DD^ was not affected by NMDA treatment (control, 100 ± 9%, *n* =14 cells; NMDA, 93± 8%; *n* = 12 cells; p = 0.53 by two-tailed Student’s t-test; Figures 4C and D). These results indicate that phosphorylation of GluA1-MPR inhibits NMDA-induced AMPAR endocytosis during LTD.

**Figure 4.**
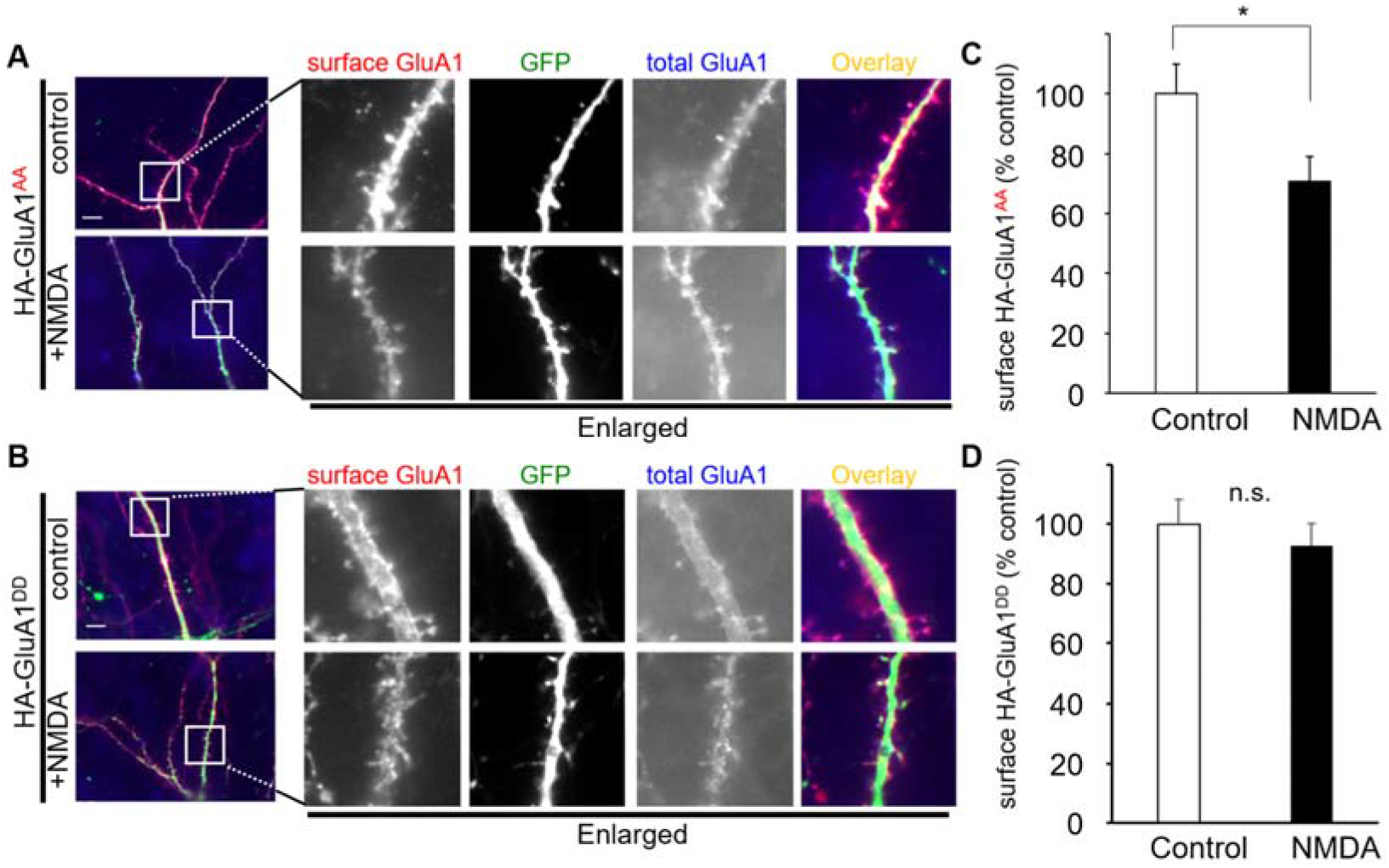
MPR regulates NMDA-induced AMPAR internalization. **A, B.** Immunocytochemical analysis of the effects of MPR on NMDA-induced trafficking of cell surface GluA1. Cultured hippocampal neurons expressing EGFP and HA-tagged dephosphomimetic GluA1 (GluA1^AA^) (**A**) or phosphomimetic GluA1 (GluA1^DD^) (**B**) were treated with 50 μM NMDA for 10 min and stained for surface HA-GluA1 (red). After treatment with Triton-X, neurons were immunostained for total HA-GluA1 (blue). The dendritic regions marked by squares were enlarged in the panels to the right. Scale bar, 10 μm. **C, D.** Quantification of NMDA-induced reduction in the ratio of the surface to total GluA1 fluorescence intensities with (NMDA) or without (control) NMDA treatment. Data are represented as the ratio of surface HA-GluA1 immunoreactivity normalized by total HA-GluA1 immunoreactivity. The ratio in control neurons was defined as 100% (*n* = 12-14). Data are presented as mean + SEM. *p < 0.05; n.s.= not significant two-tailed student’s t-test.

### Phosphorylation of GluA1-MPR regulates trafficking to the late endosome/lysosome

The number of cell-surface AMPARs is determined by the balance between endocytosis and exocytosis. To clarify the effect of phosphorylation of GluA1-MPR on AMPAR trafficking, we performed an antibody feeding assay (17) (Figure 5 A). HA-GluA1 on the cell surface of living neurons was first labeled with an anti-HA antibody and NMDA was applied to the neurons to induce AMPAR endocytosis. After removal of the anti-HA antibody remaining on the cell surface by acid treatment, the population of HA-GluA1 that was endocytosed by the NMDA treatment and recycled to the cell surface within 30 min was specifically visualized. The antibody feeding assay indicated that the amount of recycled HA-GluA1^DD^ was significantly larger than that of HA-GluA1^AA^ (HA-GluA1^AA^, 100 ± 11%, *n* = 13 cells; HA-GluA1^DD^, 127 ± 6%, *n* = 14 cells; p = 0.047 by two-tailed Student’s t-test; Figures 5B and C). These results indicate that although HA-GluA1^DD^ was endocytosed in response to NMDA treatment, it was recycled back to the cell surface.

**Figure 5.**
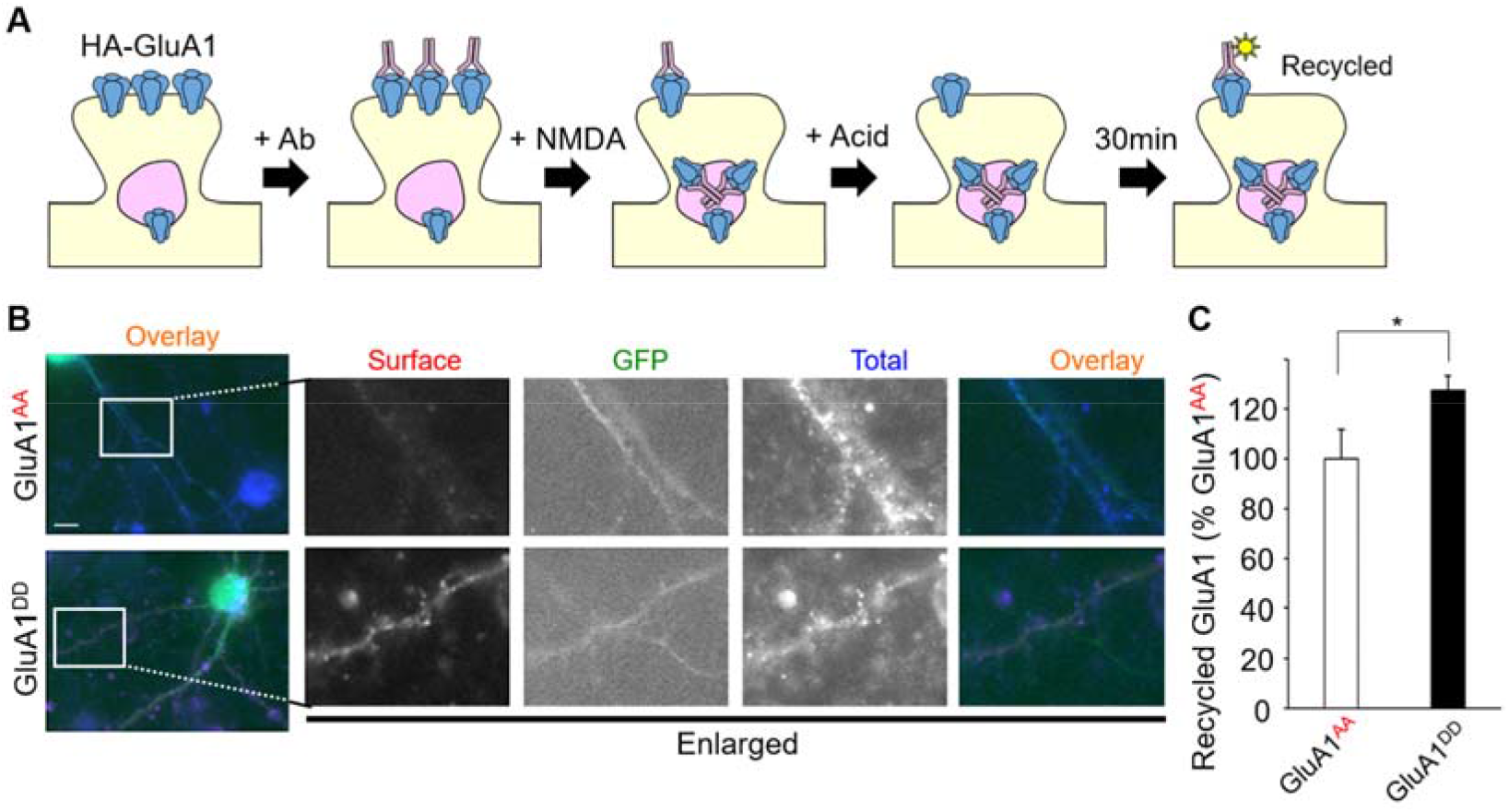
Phosphorylation of MPR increased recycling of GluA1 to the cell surface. **A.** Schematic drawing of antibody feeding assay. Living neurons expressing HA-GluA1 mutants were labeled with an anti-HA antibody. After NMDA treatment, remaining cell surface antibodies were removed by acid treatment. After a 30 min incubation to allow the recycling of HA-GluA1, neurons were fixed and recycled, and internal HA-GluA1 was visualized by Alexa546- and Alexa350-conjugated secondary antibodies, respectively. **B.** Immunocytochemical analysis of the effects of the MPR phosphorylation on the recycling of GluA1 after NMDA treatment. Cultured living hippocampal neurons expressing HA-GluA1^AA^ or HA-GluA1^DD^ were subjected to the antibody feeding assay. The dendritic regions marked by white squares are enlarged in the panels to the right. Scale bar, 10 μm. **C.** Quantification of the recycled GluA1. Data are represented as the ratio of recycled HA-GluA1 staining/total HA-GluA1 staining intensity. The ratio of HA-GluA1^AA^ was defined as 100% (*n* = 8-9 cells). Data are presented as mean + SEM. *p < 0.05 two-tailed student’s t-test.

To gain mechanistic insight into how phosphorylation of GluA1-MPR affects AMPAR trafficking, we co-expressed HA-GluA1^AA^ or HA-GluA1^DD^ with EGFP-tagged Rab4 to label early endosomes in hippocampal neurons. We also used EGFP-Rab7 to detect late endosomes or/and lysosomes and immunostained MAP2 to identify dendrites. HA-GluA1^AA^ immunoreactivity was co-localized with Rab4 at 3 min, and Rab7 at 10 min along dendrites after NMDA treatment (Figures 6A and B). In contrast, although HA-GluA1^DD^ immunoreactivity was co-localized with Rab4 at 3 min, it did not overlap with Rab7 at 10 min after NMDA treatment (Figure 6 A and B). Quantitative analysis indicated that HA-GluA1^DD^ and HA-GluA1^AA^ were similarly co-localized with Rab4 at 3 min after NMDA treatment (Figure 6 C, n = 5−6 cells, p = 0.57 by two-tailed Student’s *t*-test). In addition, HA-GluA1^DD^ showed significantly lower levels of co-localization with Rab7 than HA-GluA1^AA^ at 10 min after NMDA treatment (Figure 6 D, n = 6 cells each, p = 0.006, two-tailed Student’s *t*-test). These results indicate that phosphorylation of GluA1-MPR regulates NMDA-induced AMPAR endocytosis by controlling the transport of AMPARs from early endosomes to late endosomes/lysosomes.

**Figure 6.**
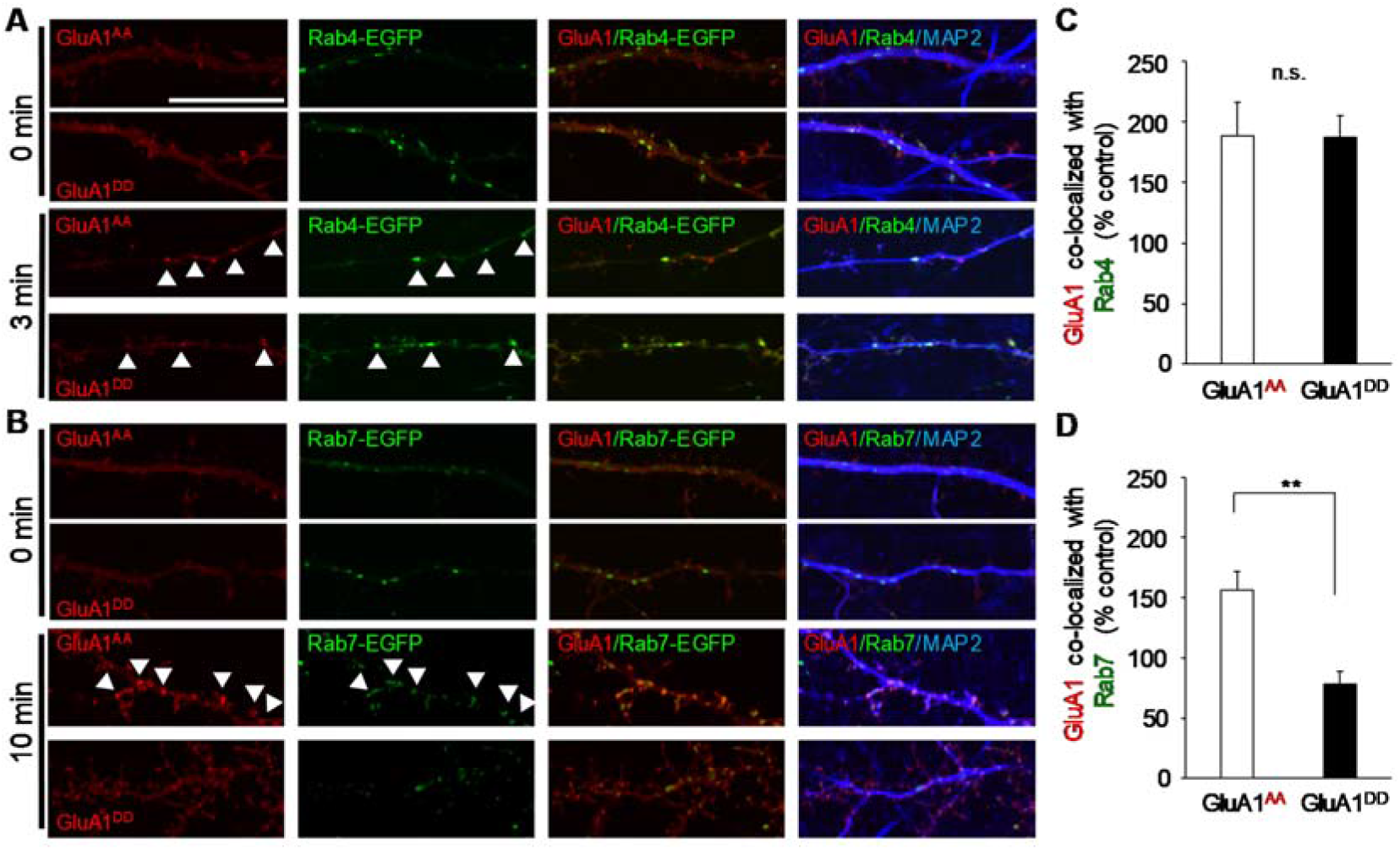
Phosphorylation of MPR inhibits the transport of GluA1 to late endosomes and lysosomes. **A.** Colocalization of HA-tagged mutant GluA1 with an early endosome marker, EGFP-tagged Rab4 at 0 min, and 3 min after NMDA treatment. Scale bar, 10 μm. **B.** Colocalization of HA-tagged mutant GluA1 with a late endosome/lysosome marker, EGFP-tagged Rab7 at 0 min, and 10 min after NMDA treatment. **C, D.** Quantification of the colocalization of HA-tagged mutant GluA1 with Rab proteins. Data are represented as the ratio of colocalized HA-GluA1 staining/total HA-GluA1 staining intensity. The ratio in the neurons without NMDA stimulation (0 min) was defined as 100% (*n* = 5-6 cells). Data are presented as mean + SEM *p < 0.05 n.s. not significant two-tailed student’s t-test.

### GluA1-MPR regulates heteromeric AMPAR trafficking

Endogenous AMPARs mainly exist as diheteromeric GluA1-GluA2 and GluA2-GluA3 receptors in the mammalian brain (20) (21). To examine whether the phosphorylation of GluA1-MPR affects the trafficking of heteromeric AMPARs composed of GluA1 and GluA2, we expressed HA-tagged GluA2 and untagged phosphomimetic GluA1^DD^ or dephosphomimetic GluA1^AA^ in cultured hippocampal neurons. After treatment with NMDA (50 μM) for 10 min, the cell surface and total GluA2 were sequentially detected by an anti-HA antibody before and after permeabilizing the plasma membrane (Figures 7A and B). The intensity of cell surface HA-GluA2 immunoreactivity was significantly reduced by the NMDA treatment in neurons co-expressing GluA1^AA^ (Figure 7 C, control, 100 ± 5%; NMDA, 81 ± 5%, n = 17 cells each; p = 0.02 by two-tailed Student’s *t*-test), but not in neurons co-expressing GluA1^DD^ (Figure 7 D, control, 100 ± 3%; NMDA, 111 ± 6%; n = 15 cells each; p = 0.08 by two-tailed Student’s *t*-test). These results indicate that the phosphorylation status at the MPR of GluA1 dominantly affects heteromeric AMPAR endocytosis during LTD.

**Figure 7.**
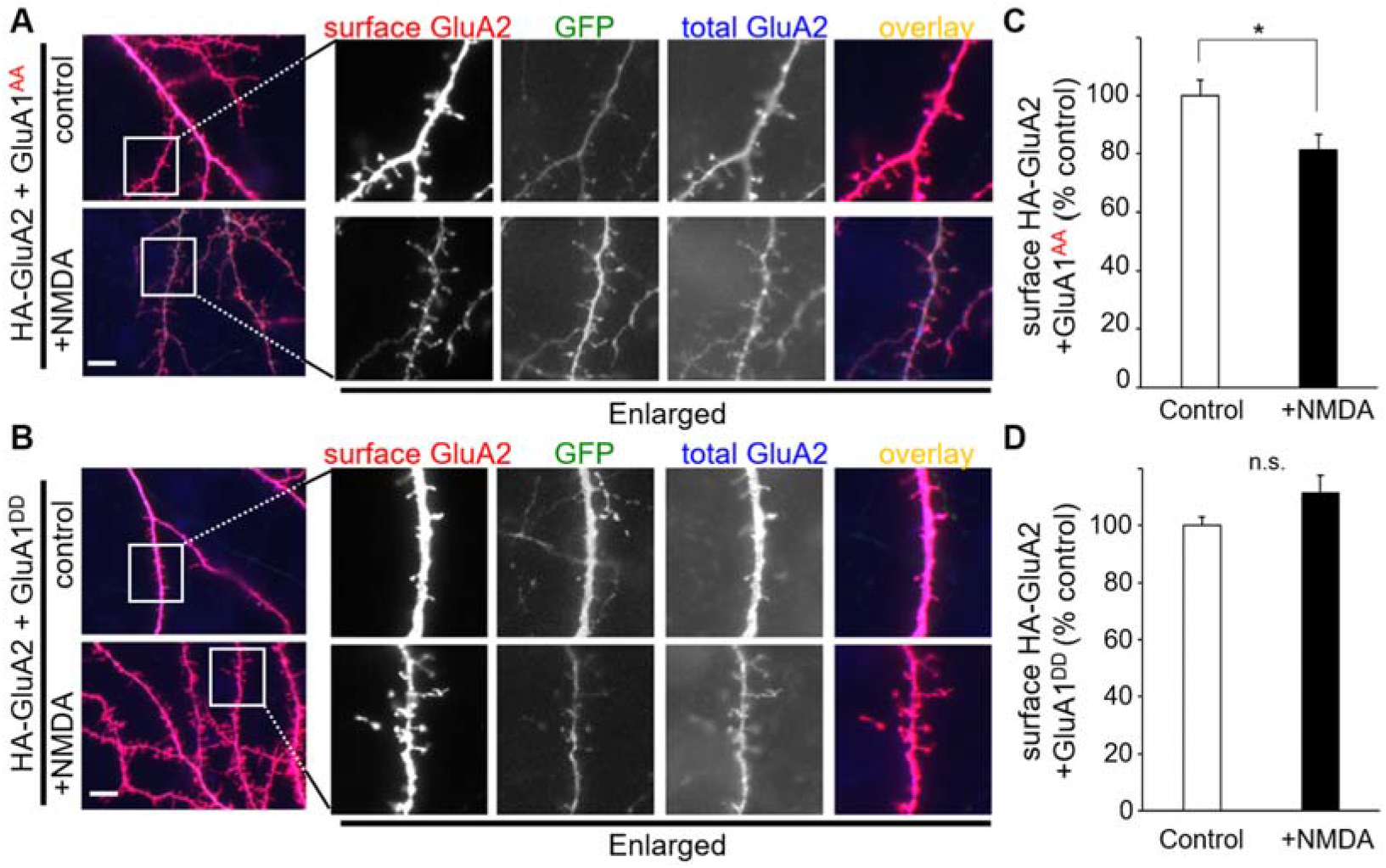
MPR regulates NMDA-induced trafficking of heteromeric AMPA receptors. **A, B.** Immunocytochemical analysis of the effects of MPR on NMDA-induced trafficking of cell surface heteromeric AMPA receptors. Cultured hippocampal neurons expressing wild-type HA-GluA2 together with GluA1^AA^ (**A**) or GluA1^DD^ (**B**) were treated with 50 μM NMDA for 10 min and stained for surface HA-GluA2 (red). After Triton X treatment, neurons were stained for total HA-GluA2 (blue). The dendritic regions marked by white squares were enlarged in the panels to the right. Scale bar, 10 μm. **C, D.** The bar graphs represent the quantification of NMDA-induced reduction in the amount of cell surface HA-GluA2 in the presence of GluA1^AA^ (**C**) or GluA1^DD^ (**D**). Data are represented as the ratio of surface HA-GluA2 staining/total HA-GluA2 staining intensity. The ratio in control neurons was defined as 100% (*n* = 15-17 cells). Data are presented as mean + SEM *p < 0.05; n.s.= not significant two-tailed student’s *t*-test.

## Discussion

It has been unclear whether and how subunit-specific rules of AMPAR trafficking are related to subunit-independent, TARP-mediated AMPAR trafficking mechanisms during LTP/LTD. In the present study, we showed that phosphomimetic mutations of GluA1-MPR inhibited AP-3 binding to STG and late endosomal/lysosomal trafficking of AMPARs, which is required for LTD expression (6,22). Thus, together with earlier findings, we propose a model in which STG-dependent and GluA1-MPR-dependent AMPAR trafficking mechanisms interact with each other during LTD in hippocampal neurons (Figure 8). At postsynaptic sites, AMPARs are stabilized by anchoring proteins, such as PSD-95, which bind to highly phosphorylated STG (16). NMDAR-induced dephosphorylation of STG releases the anchor so that the AMPAR-STG complex laterally diffuses to the endocytic zones. In the endocytic zone, AP-2 accumulates (23) and binds to dephosphorylated STG to induce clathrin-mediated endocytosis of the AMPAR-STG complex. In the early endosome, AP-2 is replaced with AP-3 to mediate transport to the late endosomes/lysosomes (Figure 8 A). When an AMPAR contains GluA1, in which the MPR remains phosphorylated, AP-3 cannot associate with STG and the AMPAR-STG complex is recycled back to the cell surface by interacting with 4.1N (10,11) (Figure 8B).

**Figure 8.**
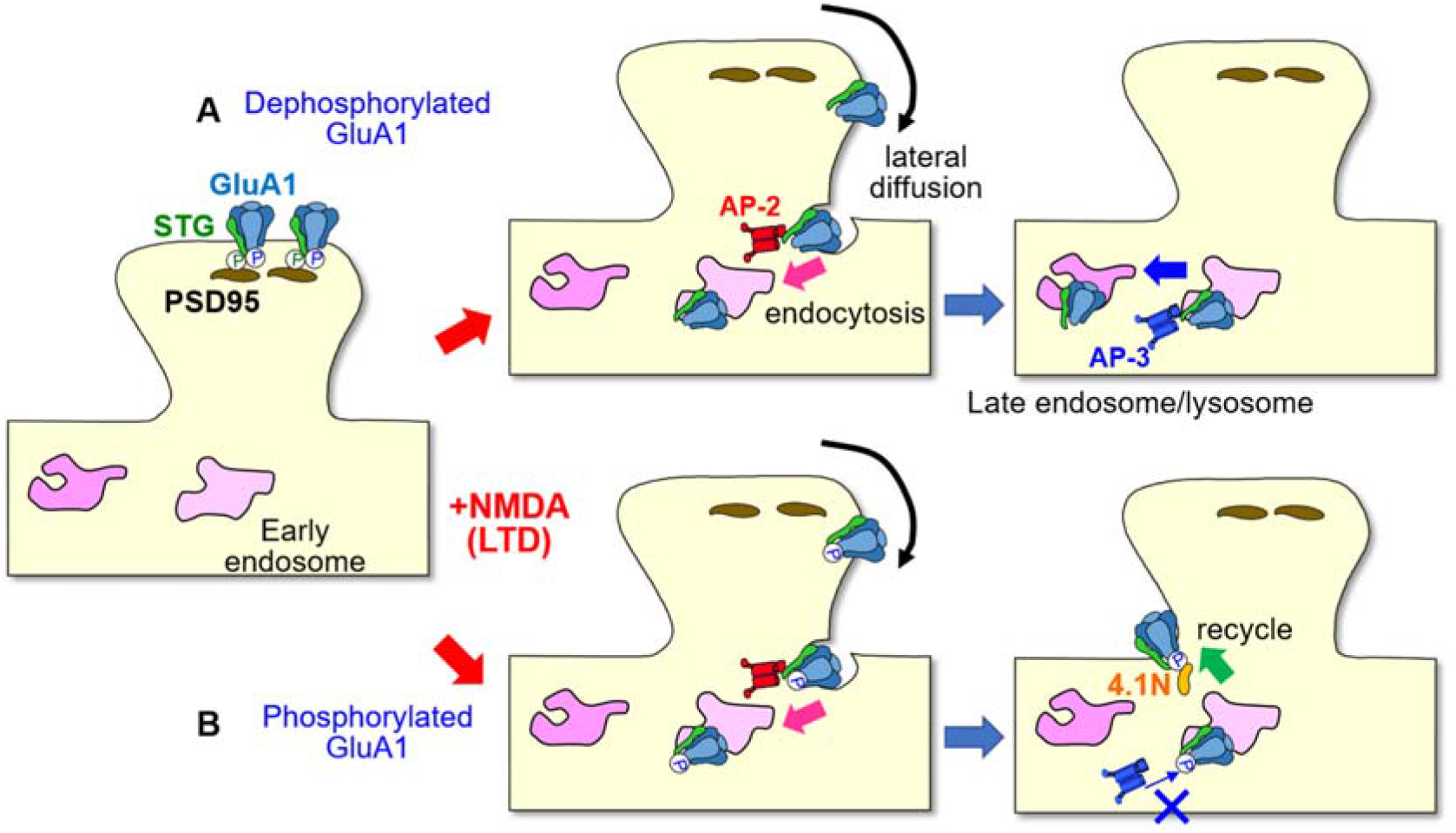
A model for AMPAR trafficking during LTD achieved by a cross-talk between subunit-dependent and independent mechanisms. An auxiliary AMPAR subunit, STG, stabilizes postsynaptic AMPARs by binding to anchoring proteins, such as PSD95. LTD-inducing stimuli dephosphorylates the C-terminus of STG and triggers lateral diffusion of the AMPAR-STG complex by reducing the binding affinity of STG to PSD-95. At the endocytic zone, dephosphorylated STG binds to AP-2 to initiate clathrin-dependent endocytosis of the AMPAR-STG complex. In the early endosomes, AP-2 is eventually replaced with AP-3 to facilitate late endosomal/lysosomal trafficking of the AMPAR-STG complex to express LTD (**A**). In contrast, AMPARs containing GluA1 behave differently depending on the phosphorylation status of the MPR, which only occurs in the GluA1 subunit. When the MPR of GluA1 remains phosphorylated, AP-3 cannot be effectively recruited to the AMPAR-STG complex. Such AMPARs are transported back to the cell surface, resulting in impaired LTD (**B**).

### Hierarchy of AMPAR trafficking mediated by GluA subunits and phosphorylation

Although AMPAR subunits and posttranslational modifications determine the types and extent of synaptic plasticity, a hierarchy may exist such that certain AMPARs are disproportionally recruited to or removed from synapses during LTP and LTD (3). This hierarchy hypothesis explains why LTP (12) and LTD (13) could still be induced in a manner independent of AMPAR subunits. However, it remains unclear how a hierarchy is determined by the subunit-dependent phosphorylation of AMPARs. We postulate that phosphorylation of GluA subunits affects two steps: anchoring at postsynaptic sites and endocytosis or exocytosis to or from plasma membranes.

For LTD, GluA2 has shown to play a major role in the hierarchy of AMPAR endocytosis in many brain regions (3). Specifically, phosphorylation of GluA2 Ser880 regulates LTD in the cerebellum (7) and the hippocampus (24). This effect is likely explained by the anchoring of GluA2-containing AMPARs by GRIP1/2 and PICK1 (25,26). Phosphorylation at Ser880 by PKC releases GluA2 from the GRIP1/2 anchor during cerebellar LTD (27,28). However, surface AMPARs are tightly associated with TARP, through which the AMPAR-TARP complex is anchored to postsynaptic sites. Thus, the release from GRIP could not fully explain the dominant role of GluA2 during LTD.

At the endocytic zone, AMPARs need to be recognized by AP-2 for clathrin-dependent endocytosis. Although the MPR of GluA2 was shown to bind to the μ2 subunit of AP-2 (29), μ2 is mainly recruited to AMPARs by binding to dephosphorylated STG in a manner independent of GluA subunits (17) and their phosphorylation status (Figure 1). GluA2, in which Ser880 is phosphorylated, could bind to PICK1 at the endocytic zone, which has been shown to recruit the α subunit of AP-2 and dynamin (30). Thus, the dominant role of GluA2 in LTD could be partly attributed to its preferential binding to PICK1.

After endocytosis, AMPARs need to be trafficked to late endosomes/lysosomes for LTD expression (6,22). Unlike μ2, the μ3 subunit of AP-3 could not be recruited to STG unless the MPR of GluA1 was fully dephosphorylated (Figure 8 B). Thus, the absence of phosphorylation sites at the MPR of GluA2 (Figure 1 A) could also contribute to the preferential role of GluA2-containing and GluA1-lacking AMPARs in LTD expression.

Phosphorylation of the MPR of GluA1 by PKC was previously shown to promote synaptic incorporation of AMPARs during LTP (10,11). Similarly, GluA1, which contained phosphomimetic mutations in the MPR, was recycled from the endosome to the cell surface (Figure 5). Because AMPARs are reported to be exocytosed from recycling endosomes (31), phosphorylation-dependent binding to μ3 by the MPR of GluA1 may also explain the subunit-selective hierarchy in LTP expression.

### Towards a unified theory of AMPAR trafficking

There remain many questions about how other phosphorylation sites of GluA subunits affect the hierarchy of AMPAR trafficking. For example, although phosphorylation at Ser845 of GluA1 is required for LTD induction (8,9), the mechanisms by which such subunit-specific phosphorylation affects LTD is achieved, remain unclear. Recently, phosphorylation at Ser845 was shown to transiently recruit GluA1-containing, Ca^2+^-permeable AMPARs, to postsynaptic sites to fully activate calcineurin during LTD (32). Indeed, calcineurin is absolutely required to dephosphorylate TARP to release the AMPAR-TARP complex from the postsynaptic anchor during hippocampal and cerebellar LTD (15,33). However, it is unclear how phosphorylation at Ser845 mediates preferential trafficking of GluA1 to postsynaptic sites. Similarly, the mechanisms by which phosphorylation at Ser831 of GluA1 contribute to LTP, remain unclear. Interestingly, phosphorylation at Ser831 and Ser845 was shown to work in concert with Ser818 phosphorylation to trigger the stable incorporation of GluA1 during hippocampal LTP (10). Thus, the effect of phosphorylation at Ser831 and Ser845 on AMPAR trafficking could be partly attributed to phosphorylation levels at the MPR, which determine the association with AP-3 and 4.1N. Because differential phosphorylation of AMPARs is reported in certain mouse models of neuropsychiatric disease, such as fragile X mental retardation (34), further studies are warranted to clarify the molecular mechanisms by which phosphorylation and other posttranslational modifications regulate the hierarchy of AMPAR trafficking.

## Experimental procedures

### Chemicals and Antibodies

NMDA was purchased from Tocris Bioscience. Commercial antibodies: anti-GluA1 (Millipore 04-855), anti-stargazin (Sigma C8206), anti-GST (Amersham RPN1236), anti-HA, (Covance 901501), anti-FLAG (Sigma F7425), and anti-MAP-2 (Millipore AB5622) antibodies; Alexa-350 (Thermo Fisher A-11045), 405 (Thermo Fisher A-31556), 488 (Thermo Fisher A-11008), 546 (Thermo Fisher A-11003), HRP (Rockland 18-8816-33, 18-8817-33) conjugated secondary antibodies.

### Construction and Transfection or Transformation of Expression Plasmids

Using a PCR method and Pyrobest (Takara), the serine residues encoding S816, S818, S831, and S845 in mouse GluA1 cDNA were mutated to encode aspartate or alanine. The cDNA encoding HA was added to the 5’ end (immediately following the signal sequence) of mutant GluA1 and wild-type GluA2. The cDNA encoding flag-tag was added to the 3’ end (immediately upstream of the stop codon) of mouse μ2 or mouse μ3A cDNAs. The nucleotide sequences of the amplified open reading frames were confirmed by bidirectional sequencing. After the cDNAs were cloned into the expression vectors, either pTracer (Invitrogen) or pCAGGS (provided by Dr. J Miyazaki, Osaka University, Osaka, Japan), the constructs were transfected into human embryonic kidney 293T (HEK293T) cells using the Ca^2+^-phosphate method or were transfected into cultured hippocampal neurons using Lipofectamine 2000 (Invitrogen).

For the expression of GST-fusion protein, the cDNA encoding the C-terminal region of wild-type or mutant TARPs was amplified by PCR and cloned into pGEX 4T-2. *E. coli*. BL21(DE3) was transformed by pGEX expression vectors and the expression of GST fusion proteins was induced by the addition of 0.1mM IPTG. BL21(DE3) was disrupted by sonication in phosphate-buffered saline (PBS), and GST fusion proteins were purified using the Glutathione Sepharose column (Amersham Pharmacia).

### Culture of Hippocampal Neuron

Hippocampi dissected from E16/17 ICR mice were treated with 10 U ml^−1^ trips and 100 U ml^−1^ DNase in Dulbecco’s modified Eagle’s medium at 37°C for 20 min. The dissociated hippocampal neurons were plated on polyethyleneimine (PEI)-coated glass coverslips and cultured in Neurobasal medium (Invitrogen) with B-27 (Gibco) or NS21 supplement (35) and 0.5 mM L-glutamine. After 7-10 DIV culture, neurons were transiently transfected with plasmids using Lipofectamine 2000, and used for the AMPA receptor endocytosis or recycling assays.

### Assay for AMPA receptor Endocytosis

Hippocampal neurons transfected with pCAGGS expression vectors for mutant HA-GluA1 plus GFP or wild-type HA-GluA2 plus mutant GluA1 were stimulated with 50 μM NMDA for 10 min and fixed in 4% paraformaldehyde without permeabilization, for 10 min at room temperature. After fixed neurons were washed with PBS and incubated with a blocking solution (2% BSA and 2% normal goat serum in PBS), surface HA-GluA1 or HA-GluA2 were labeled with the anti-HA antibody (1:1000) and visualized with Alexa 546-secondary antibody (1:1000). To label total HA-GluA1 or HA-GluA2, neurons were permeabilized and blocked with a blocking solution containing 0.4% Triton X-100, and incubated with the anti-HA antibody (1:1000) and Alexa 350-secondary antibodies (1:1000). Fluorescence images were captured using a fluorescence microscope (BX60, Olympus) equipped with a CCD camera (DP 70, Olympus) and analyzed using IP-Lab software (Scanalytics). For statistical analysis of the surface expression level of HA-GluA1 or HA-GluA2, the intensity of Alexa 546 for surface HA-GluA1 or HA-GluA2 was measured and normalized using the intensity of Alexa 350 for total HA-GluA1 or HA-GluA2. The fluorescence intensity on the dendrites at least 20 μm away from the soma, was measured.

### Assay for AMPA receptor recycling

Recycling of AMPA receptors was analyzed by the method described by Nooh et al. (36). Living hippocampal neurons transfected with plasmids for mutant HA-GluA1 were labeled with anti-HA antibody (1:100) for 1 h. After washing out the excess amount of antibody, neurons were stimulated with 50 μM NMDA for 3 min. After washing out the NMDA, neurons were treated with 0.5 M NaCl and 0.2 M acetic acid for 4 min at 0°C. After washing out NaCl and acetic acid, neurons were incubated for 30 min at 37°C in a neurobasal medium with B27 supplement. The neurons were then fixed in 4% paraformaldehyde without permeabilization, for 10 min at room temperature. After fixed neurons were washed with PBS and incubated in a blocking solution (2% BSA and 2% normal goat serum in PBS), surface HA antibody was visualized with Alexa 546-secondary antibody (1:1000). To label internalized HA-GluA1, neurons were permeabilized and blocked with the blocking solution containing 0.4% Triton X-100 and incubated with the Alexa 350-secondary antibodies (1:1000). Fluorescence images were captured by a fluorescence microscope equipped with a CCD camera and analyzed using IP-Lab software. For statistical analysis of the recycled HA-GluA1, the intensity of Alexa 546 for recycled HA-GluA1 was measured and normalized using the intensity of Alexa 350 for internalized HA-tagged GluA1. The fluorescence intensity on the dendrites at least 20 μm away from the soma, was measured.

### Colocalization assay of HA-GluA1 and Rab proteins

Hippocampal neurons transfected with pCAGGS expression vectors for mutant HA-GluA1, Rab4, or Rab7-EGFP were stimulated with 50 μM NMDA for 3 or 10 min and fixed in 4% paraformaldehyde. After fixed neurons were washed with PBS and incubated with a blocking solution (2% BSA and 2% normal goat serum 0.4% Triton-X in PBS), the neurons were incubated with the anti-HA antibody (1:1000) and anti-MAP-2 antibody (1:1000) for 1 h at room temperature. After washing with PBS, neurons were incubated with Alexa 546- and Alexa 405-secondary antibodies (1:1000; Invitrogen). Fluorescence images were captured using a confocal microscope (FV1200, Olympus) and analyzed using IP-Lab software (Scanalytics). To statistically analyze the colocalization of the HA-GluA1 and Rab proteins, the intensities of Alexa 546 on the EGFP-positive regions were measured and normalized using the total intensity of Alexa 546. The fluorescence intensity on the dendrites at least 20 μm away from the soma, was measured.

### Immunoprecipitation, Pull-Down Assay, and Immunoblot Assays

Transfected HEK293T cells were solubilized in 500 μL of TNE buffer (50 mM NaCl, 10 % NP-40, 20 mM EDTA, 0.1 % SDS, 50 mM Tris-HCl, pH 8.0) supplemented with a protease inhibitor cocktail (Calbiochem).

For the immunoprecipitation assays, 5 μl of anti-GluA1 (Millipore) was added to the samples, and the mixture was incubated for 1 h at 4°C. Then, 50 μL of protein G-conjugated agarose (Amersham) was added, and this mixture was incubated for 1 h at 4°C. After the precipitates were washed four times with 500 μL of TNE buffer, 50 μL of SDS-PAGE sample buffer was added and the samples were incubated for 5 min at 95°C. After centrifugation, 5 μL of the supernatant was analyzed using immunoblotting with anti-Flag (Sigma) or other primary antibodies, TrueBlot HRP-conjugated secondary antibody (Rockland), and the Immobilon Western kit (Millipore). The chemiluminescence signals were detected by Luminograph II (ATTO) and quantified using CS Analyzer software (ATTO).

For GST pull-down assays, purified GST fusion proteins with a TARP C-terminus were incubated with the lysate of HEK293T cells expressing the μ subunit of adaptor protein in the presence or absence of 500μM of peptides corresponding to the MPR of AMPA receptors. After a 1 h incubation at 4°C, GST proteins were pulled down by glutathione sepharose resins (Amersham), and the precipitates were analyzed by immunoblot analysis.

For the biotinylated peptide pull-down assay, purified GST fusion proteins with a TARP C-terminus were incubated with the biotinylated peptide corresponding to the MPR of AMPA receptors. After a 1 h incubation at 4°C, biotinylated peptides were pulled down using streptavidin-conjugated magnetic beads (Invitrogen), and the precipitates were analyzed by immunoblot analysis

## Data availability

All data described are presented either within the article or in the supporting information.

## Acknowledgments

This work was supported by the CREST from JST (S.M. and M.Y.), MEXT (16H06461,15H05772 to M.Y.; 17K07048 to S.M.). We thank A. Takahashi, J. Motohashi, and S. Narumi for their technical assistance.

## Author contribution

S. M. performed the experiments and analyzed the data. S.M. and M.Y. conceived and coordinated the study and wrote the paper. All authors reviewed the results and approved the final version of the manuscript.

## Abbreviations

The abbreviations used are:

LTP: long term potentiation
LTD: long term depression
AMPA: α-Amino-3-hydroxy-5-methyl-4-isoxazolepropionic Acid
MPR: membrane-proximal region
STG: stargazing
NMDA: N-methyl-D-aspartate
C-terminus, CaMKII: calcium/calmodulin-dependent protein kinase II
PKA: protein kinase A
PKC: protein kinase C
TARP: transmembrane AMPAR regulatory protein
PSD95: postsynaptic density 95
HEK293: human embryonic kidney 293
GST: glutathione S-transferase
HA: hemagglutinin
MAP2: Microtubule-associated protein 2
PBS: phosphate-buffered saline
BSA: bovine serum albumin

## REFERENCES

1. Collingridge, G. L., Peineau, S., Howland, J. G., and Wang, Y. T. (2010) Long-term depression in the CNS. Nat Rev Neurosci 11, 459–473 doi: 10.1038/nrn2867

2. Nicoll, R. A. (2017) A Brief History of Long-Term Potentiation. Neuron 93, 281–290 doi: 10.1016/j.neuron.2016.12.015

3. Diering, G. H., and Huganir, R. L. (2018) The AMPA Receptor Code of Synaptic Plasticity. Neuron 100, 314–329 doi: 10.1016/j.neuron.2018.10.018

4. Groc, L., and Choquet, D. (2020) Linking glutamate receptor movements and synapse function. Science 368 doi: 10.1126/science.aay4631

5. Shi, S., Hayashi, Y., Esteban, J. A., and Malinow, R. (2001) Subunit-specific rules governing AMPA receptor trafficking to synapses in hippocampal pyramidal neurons. Cell 105, 331–343 doi: 10.1016/s0092-8674(01)00321-x

6. Lee, S. H., Simonetta, A., and Sheng, M. (2004) Subunit rules governing the sorting of internalized AMPA receptors in hippocampal neurons. Neuron 43, 221–236 doi: 10.1016/j.neuron.2004.06.015

7. Chung, H. J., Steinberg, J. P., Huganir, R. L., and Linden, D. J. (2003) Requirement of AMPA receptor GluR2 phosphorylation for cerebellar long-term depression. Science 300, 1751–1755 doi: 10.1126/science.1082915

8. Lee, H. K., Barbarosie, M., Kameyama, K., Bear, M. F., and Huganir, R. L. (2000) Regulation of distinct AMPA receptor phosphorylation sites during bidirectional synaptic plasticity. Nature 405, 955–959 doi: 10.1038/35016089

9. Lee, H. K., Takamiya, K., Han, J. S., Man, H., Kim, C. H., Rumbaugh, G., Yu, S., Ding, L., He, C., Petralia, R. S., Wenthold, R. J., Gallagher, M., and Huganir, R. L. (2003) Phosphorylation of the AMPA receptor GluR1 subunit is required for synaptic plasticity and retention of spatial memory. Cell 112, 631–643 doi: 10.1016/s0092-8674(03)00122-3

10. Boehm, J., Kang, M. G., Johnson, R. C., Esteban, J., Huganir, R. L., and Malinow, R. (2006) Synaptic incorporation of AMPA receptors during LTP is controlled by a PKC phosphorylation site on GluR1. Neuron 51, 213–225 doi: 10.1016/j.neuron.2006.06.013

11. Lin, D. T., Makino, Y., Sharma, K., Hayashi, T., Neve, R., Takamiya, K., and Huganir, R. L. (2009) Regulation of AMPA receptor extrasynaptic insertion by 4.1N, phosphorylation and palmitoylation. Nat Neurosci 12, 879–887 doi: 10.1038/nn.2351.

12. Granger, A. J., Shi, Y., Lu, W., Cerpas, M., and Nicoll, R. A. (2013) LTP requires a reserve pool of glutamate receptors independent of subunit type. Nature 493, 495–500 doi: 10.1038/nature11775

13. Granger, A. J., and Nicoll, R. A. (2014) LTD expression is independent of glutamate receptor subtype. Front Synaptic Neurosci 6, 15 doi: 10.3389/fnsyn.2014.00015

14. Nicoll, R. A., Tomita, S., and Bredt, D. S. (2006) Auxiliary subunits assist AMPA-type glutamate receptors. Science 311, 1253–1256 doi: 10.1126/science.1123339

15. Tomita, S., Stein, V., Stocker, T. J., Nicoll, R. A., and Bredt, D. S. (2005) Bidirectional synaptic plasticity regulated by phosphorylation of stargazin-like TARPs. Neuron 45, 269–277 doi: 10.1016/j.neuron.2005.01.009

16. Sumioka, A., Yan, D., and Tomita, S. (2010) TARP phosphorylation regulates synaptic AMPA receptors through lipid bilayers. Neuron 66, 755–767 doi: 10.1016/j.neuron.2010.04.035.

17. Matsuda, S., Kakegawa, W., Budisantoso, T., Nomura, T., Kohda, K., and Yuzaki, M. (2013) Stargazin regulates AMPA receptor trafficking through adaptor protein complexes during long-term depression. Nat Commun 4, 2759 doi: 10.1038/ncomms3759

18. Zhou, Z., Liu, A., Xia, S., Leung, C., Qi, J., Meng, Y., Xie, W., Park, P., Collingridge, G. L., and Jia, Z. (2018) The C-terminal tails of endogenous GluA1 and GluA2 differentially contribute to hippocampal synaptic plasticity and learning. Nat Neurosci 21, 50–62 doi: 10.1038/s41593-017-0030-z

19. Lu, W., and Roche, K. W. (2012) Posttranslational regulation of AMPA receptor trafficking and function. Curr Opin Neurobiol 22, 470–479 doi: 10.1016/j.conb.2011.09.008

20. Lu, W., Shi, Y., Jackson, A. C., Bjorgan, K., During, M. J., Sprengel, R., Seeburg, P. H., and Nicoll, R. A. (2009) Subunit composition of synaptic AMPA receptors revealed by a single-cell genetic approach. Neuron 62, 254–268 doi: 10.1016/j.neuron.2009.02.027

21. Zhao, Y., Chen, S., Swensen, A. C., Qian, W. J., and Gouaux, E. (2019) Architecture and subunit arrangement of native AMPA receptors elucidated by cryo-EM. Science 364, 355–362 doi: 10.1126/science.aaw8250

22. Fernandez-Monreal, M., Brown, T. C., Royo, M., and Esteban, J. A. (2012) The balance between receptor recycling and trafficking toward lysosomes determines synaptic strength during long-term depression. J Neurosci 32, 13200–13205 doi: 10.1523/JNEUROSCI.0061-12.2012.

23. Unoki, T., Matsuda, S., Kakegawa, W., Van, N. T., Kohda, K., Suzuki, A., Funakoshi, Y., Hasegawa, H., Yuzaki, M., and Kanaho, Y. (2012) NMDA receptor-mediated PIP5K activation to produce PI(4,5)P(2) is essential for AMPA receptor endocytosis during LTD. Neuron 73, 135–148 doi: 10.1016/j.neuron.2011.09.034

24. Seidenman, K. J., Steinberg, J. P., Huganir, R., and Malinow, R. (2003) Glutamate receptor subunit 2 Serine 880 phosphorylation modulates synaptic transmission and mediates plasticity in CA1 pyramidal cells. J Neurosci 23, 9220–9228 doi: 10.1523/JNEUROSCI.23-27-09220.2003

25. Steinberg, J. P., Takamiya, K., Shen, Y., Xia, J., Rubio, M. E., Yu, S., Jin, W., Thomas, G. M., Linden, D. J., and Huganir, R. L. (2006) Targeted in vivo mutations of the AMPA receptor subunit GluR2 and its interacting protein PICK1 eliminate cerebellar long-term depression. Neuron 49, 845–860 doi: 10.1016/j.neuron.2006.02.025.

26. Takamiya, K., Mao, L., Huganir, R. L., and Linden, D. J. (2008) The glutamate receptor-interacting protein family of GluR2-binding proteins is required for long-term synaptic depression expression in cerebellar Purkinje cells. J Neurosci 28, 5752–5755 doi: 10.1523/JNEUROSCI.0654-08.2008.

27. Matsuda, S., Launey, T., Mikawa, S., and Hirai, H. (2000) Disruption of AMPA receptor GluR2 clusters following long-term depression induction in cerebellar Purkinje neurons. EMBO J 19, 2765–2774 doi: 10.1093/emboj/19.12.2765

28. Xia, J., Chung, H. J., Wihler, C., Huganir, R. L., and Linden, D. J. (2000) Cerebellar long-term depression requires PKC-regulated interactions between GluR2/3 and PDZ domain-containing proteins. Neuron 28, 499–510 doi: 10.1016/s0896-6273(00)00128-8

29. Kastning, K., Kukhtina, V., Kittler, J. T., Chen, G., Pechstein, A., Enders, S., Lee, S. H., Sheng, M., Yan, Z., and Haucke, V. (2007) Molecular determinants for the interaction between AMPA receptors and the clathrin adaptor complex AP-2. Proc Natl Acad Sci U S A 104, 2991–2996 doi: 10.1073/pnas.0611170104

30. Fiuza, M., Rostosky, C. M., Parkinson, G. T., Bygrave, A. M., Halemani, N., Baptista, M., Milosevic, I., and Hanley, J. G. (2017) PICK1 regulates AMPA receptor endocytosis via direct interactions with AP2 alpha-appendage and dynamin. J Cell Biol 216, 3323–3338 doi: 10.1083/jcb.201701034

31. Park, M., Penick, E. C., Edwards, J. G., Kauer, J. A., and Ehlers, M. D. (2004) Recycling endosomes supply AMPA receptors for LTP. Science 305, 1972–1975 doi: 10.1126/science.1102026.

32. Sanderson, J. L., Gorski, J. A., and Dell’Acqua, M. L. (2016) NMDA Receptor-Dependent LTD Requires Transient Synaptic Incorporation of Ca(2)(+)-Permeable AMPARs Mediated by AKAP150-Anchored PKA and Calcineurin. Neuron 89, 1000–1015 doi: 10.1016/j.neuron.2016.01.043.

33. Nomura, T., Kakegawa, W., Matsuda, S., Kohda, K., Nishiyama, J., Takahashi, T., and Yuzaki, M. (2012) Cerebellar long-term depression requires dephosphorylation of TARP in Purkinje cells. Eur J Neurosci 35, 402–410 doi: 10.1111/j.1460-9568.201107963.x.

34. Tian, M., Zeng, Y., Hu, Y., Yuan, X., Liu, S., Li, J., Lu, P., Sun, Y., Gao, L., Fu, D., Li, Y., Wang, S., and McClintock, S. M. (2015) 7, 8-Dihydroxyflavone induces synapse expression of AMPA GluA1 and ameliorates cognitive and spine abnormalities in a mouse model of fragile X syndrome. Neuropharmacology 89, 43–53 doi: 10.1016/j.neuropharm.2014.09.006.

35. Chen, Y., Stevens, B., Chang, J., Milbrandt, J., Barres, B. A., and Hell, J. W. (2008) NS21: re-defined and modified supplement B27 for neuronal cultures. J Neurosci Methods 171, 239–247 doi: 10.1016/j.jneumeth.2008.03.013.

36. Nooh, M. M., Chumpia, M. M., Hamilton, T. B., and Bahouth, S. W. (2014) Sorting of beta1-adrenergic receptors is mediated by pathways that are either dependent on or independent of type I PDZ, protein kinase A (PKA), and SAP97. J Biol Chem 289, 2277–2294 doi: 10.1074/jbc.M113.513481

